# Wheat and barley stripe rust pan-genome facilitates discovery of the predominant North American lineage’s origin in somatic hybridization

**DOI:** 10.1101/2025.10.24.684071

**Authors:** Samuel Holden, Meng Li, Mehrdad Abbasi, Ramandeep Bamrah, Sang Hu Kim, Sean Formby, Janice Bamforth, Sean Walkowiak, Andrew Friskop, Upinder S. Gill, Brent McCallum, Harpinder S. Randhawa, Hadley R. Kutcher, Valentyna Klymiuk, Guus Bakkeren, Curtis Pozniak, Gurcharn S. Brar

**Affiliations:** Faculty of Agricultural, Life and Environmental Science, University of Alberta, Edmonton, AB, Canada; Faculty of Land and Food Systems, The University of British Columbia, Vancouver, BC, Canada; Agriculture and Agri Food Canada, Summerland Research and Development Centre, Summerland, BC, Canada; Faculty of Science, The University of British Columbia, Vancouver, BC, Canada; Canadian Grain Commission, Winnipeg, MB, Canada; Department of Plant Pathology, North Dakota State University, Fargo, ND, USA; Agriculture and Agri Food Canada, Morden Research and Development Centre, Morden, MB, Canada; Agriculture and Agri Food Canada, Lethbridge Research and Development Centre, Lethbridge, AB, Canada; Crop Development Centre, University of Saskatchewan, Saskatoon, SK, Canada; University of Guelph, Guelph, ON, Canada

## Abstract

Wheat and barley stripe rust, caused by *Puccinia striiformis* formae speciales *tritici* (*Pst*) and *hordei* (*Psh*) respectively is a perennial and endemic threat to those crops worldwide. Outside the Himalayan region the fungus has not been observed reproducing sexually, despite being characterised extensively over the past 100 years by researchers in many countries. Scientific literature divides *P. striiformis* into lineages termed PstS0, PstS1, up to PstS18. Here we describe the first North American *P. striiformis* pan-genome, comprising 24 *Pst* and 3 *Psh* genomes from North America including five phased assemblies using chromosome conformation capture to separate two haploid genomes. We identify the origin of the predominant North American *Pst* lineage; PstS18, as a somatic hybridization event between PstS1 and another unknown lineage. As well as identifying gene families which distinguish haplotypes on a trans-lineage and trans-species level, we also show ongoing gene family loss and expansion in clonal lineages after their emergence.

## Introduction

Pathogenic variability in rust fungi is thought to arise from random mutation, sexual recombination, and somatic hybridization [1]. In Canada and several other countries *Puccinia striiformis* f. sp *tritici (Pst)* samples are routinely collected and monitored for biosecurity and research purposes, sexual recombination is not observed, and mutation is believed to be the primary source of ongoing adaptation and evolution. Emerging races typically differ from an existing race by gain or loss (mostly former) of virulence on single host resistance gene, leading to clonal lineages of closely related races [2,3]. Although natural mutation provides ample resources for variation in the rust pathogens, it naturally produces only stepwise changes to limited numbers of genes in each round of reproduction. Recombination provides a mechanism for large-scale introduction of new genetic material as well as reshuffling of existing genes and consequent variation. Epidemic-causing lineages such as Pst7/Warrior and the stem rust pathovar Ug99 are the result of genetic novelty emerging which local crops are unprepared for, and genome sequencing has shown that novelty arose through recombination in these specific cases [4,5]. It is now well-established that recombination in rust species can be sexual, parasexual, or through somatic hybridization, although it is unclear if all species can reproduce using all mechanisms and until now there has never been genomic evidence for somatic hybridisation in *Pst* or *Psh*. Sexual recombination involves the development of the haploid fungal state on the alternate host and reproduction with a compatible mate. For *Puccinia graminis* and *Puccina pseudostriiformis* the alternate hosts are susceptible *Berberris spp.* (the barberry bush), and the same is believed to be true for *Puccinia striiformis*, although sexual reproduction has not been recorded in the wild. Sexual recombination is regularly observed in other cereal rusts such as oat crown rust (*Puccinia coronata* f. sp. *avenae*, *Pca*), stem rust (*Puccinia graminis, Pg*), and in many non-cereal rusts such as in white pine blister rust (*Cronartium ribicola*). Somatic hybridization has not been directly observed in these species but is believed to involve the fusion of dikaryotic vegetative hyphae from two colonies on any host plant, leading to an intermediate cell with four nuclei. This cell then loses two nuclei and can produce new daughter hyphae that inherit one nuclear genome intact from each of the two original colonies and can go on to develop normally and eventually form fruiting bodies. Parasexual recombination represents any intermediate state between sexual and somatic hybridization and is both rare and understudied but is inferred as an explanation for situations where nuclear exchange or crossing over occurs without going through every stage of sexual reproduction as described above.

Prior to the genomics era, detecting and confirming naturally occurring somatic hybrids in cereal rust fungi was highly challenging due to the absence of genetic markers that could resolve to the level of an individual nucleus. Instead, pre-genomics studies on the stripe rust pathogen *Pst* by many researchers [6–8] have provided robust evidence of somatic hybridization under controlled conditions. In these studies, new virulences were observed following the infection of a susceptible wheat plant with two pathogenically distinct parental races and from this evidence, somatic hybridisation in nature is inferred to be plausible, if uncommon. Now, chromosome conformation capture technology permits the matter to be empirically evaluated by comparing the complete nuclear genomes of isolates. If there is complete agreement between a single haplotype of two isolates, with no evidence of recombination, then somatic hybridisation is the only known biological explanation for the observed phenomenon. While identification of a third isolate representing the other parent is ideal, it is not necessary for the conclusion to be drawn.

The *Pst* population in North America is believed to be entirely asexual and descended from European and African isolates that established themselves following either long-distance spore dispersal across the ocean or, more likely, introduction mediated by human travel [9]. The susceptible alternate host of *Pst* does exist in parts of North America but Wang et al. [10] demonstrated that barberry does not function as an alternate host for *Pst* in the US Pacific Northwest (PNW) due to teliospore degradation and barberry phenology. From 2000 to about 2016, the *Pst* population in North America was dominated by the invasive and warmer-temperature adapted lineage PstS1, which is very similar to contemporary east African PstS1 isolates and appears to be descended from a single founding isolate [11,12]. Studies on Canadian stripe rust populations provided clear evidence of a shift toward increased virulence since 2014 [13–15]. A comprehensive population genetic study of *Pst* isolates in Canada by Brar et al. [12] was the first to identify and classify Canadian *Pst* lineages into a global context; this study identified four circulating lineages: PstS0 (old Northwestern European), PstS1 (post-2000 introduction related to east Africa), PstS18 (then named PstS1-related), and PstPr (an isolate similar to Chinese isolates of no placed lineage, with indicators of recent sexual recombination). Of the four lineages, PstS18 and PtsPr are novel, produce abundant telia and have not been reported from any continent other than North America. Based on available evidence, Brar et al.[12] hypothesized that PstS18 is a somatic hybrid and PstPr is a foreign introduction that was subsequently outcompeted as it has never been reported in subsequent surveys. Our most recent study [16] reported that PstS18 has largely replaced the PstS1 lineage in North America since 2016 and has become the predominant lineage, likely due to adaptions favouring increased sporulation in the warmer environmental conditions of western Canada, as well as spreading eastwards at least as far as Saskatchewan. However, an important biological questions of whether PstS18 is genetically distinct from PstS1 as well as where and how it emerged remained unresolved.

At the time of writing a phased genome sequence exists for *Pst* [17–19], *Pca* [20,21], and *Puccinia triticina* [22]. In this study we assemble four additional phased *Pst* genomes (Yr2016-44, T030, W047-1, W056), the first phased *Psh* genome (NDPSH), as well as 20 *Pst* and two *Psh* genome assemblies, which allowed us to develop a haplotype-resolved North American *Puccinia striiformis* pan-genome including all known North American lineages. We also included four global reference genomes. We use these data to demonstrate that the dominant North American *Pst* lineage, PstS18, is indeed the product of recent somatic hybridisation. In addition to a somatic hybridization event in *Pst*, we also identify two lineages of *Psh* (PshS1 and PshS2) and report their shared single haplotype, making this the first report of somatic hybridisation in *Psh.* Finally, we explore the extent of intra-haplotype diversity within these clonal *P. striiformis* lineages and found that each exhibits haplotype-specific gene family expansions and contractions that are shared across clonal lineages but not within them.

## Results

### Inter-lineage diversity in the *Puccinia striiformis* pan-genome is revealed by comparing phased haplotypes

We performed de-novo PacBio whole genome sequencing of 24 isolates of *Pst* and three isolates of *Psh* (**Table 1**); 23 of which were collected between 2010 and 2020, and four historical isolates (collected prior to 2010). We assembled five of these genomes using Hi-C data allowing two phased nuclear genomes to be obtained for isolates: NDPSH (*Psh*), W047-1 (PstS1), Yr2016-44 (PstS18), T030 (PstS18), and W056 (PstPr). We also included in our analysis publicly available phased reference genomes from around the world: AZ2 (Chinese isolate, F1 recombinant) [19], Pst134 E16 A+ 17+ 33+ (PstS1, Australia) [18], and Pst130 (PstS1, North American) [23] as well as DK0911 (PstS7/Warrior, Europe) [5]. Genomes were annotated for repeats and for genes prediction and functional annotation. Chromosomes were scaffolded using Pst134 E16 A+ as a reference. To make it easier to describe groups of haploid genomes with high nuclear identity from different lineages, we named our haplotype groups A-J, where members of a haplotype are more similar to one another than to any member of another haplotype (generally <1 SNP/kbp) Diploid genomes were between 149 Mbp and 205 Mbp, with a median of 153 Mbp. Predicted diploid gene counts were between 29496 and 37053, with a median of 30389 (**Table 1**).

**Table 1.** Genomes assembled in this study, and important summary statistics relating to them. Phased and pseudo-phased genomes are divided into primary (1) and alternate (2). In all cases genomes are described pre-RagTag scaffolding, as scaffolding simply joined together appropriate contigs separated by 100 N nucleotides.

We first wanted to assess the overall divergence between our samples, specifically between lineages of *Pst* and between *Pst* and *Psh* in order to evaluate *Psh* as an outgroup for *Pst*, and to examine the diversity between and within *Pst* lineages. For this, we compared gene orthologue presence/absence, coverage of each reference genome when aligned to each other genome, and SNPs/kbp across our reference genomes. To our surprise, *Psh* did not exhibit an extreme degree of difference to the *Pst* samples but was similar in magnitude to the highest level of inter-lineage *Pst* divergence: observed between AZ2 and other samples (**Supplementary File S1)**

As expected, results from comparing individual haplotypes were rather different to comparing entire genomes. Firstly, when comparing both haplotypes of the same individual, much higher levels of difference were observed versus comparisons across genomes. These values were nearly all over 5 SNP/Kbp. An exception was AZ2, which was reported specifically in Wang et al. [19] to be the result of a sexual hybridization event and to exhibit low levels of intra-specific diversity **(Supplementary File S1)**.

Secondly, comparisons of individual haplotypes showed that each isolate’s two divergent haplotypes fall into one of two ancestral groups in which all members of a group are more similar to one another than the complementary haploid genome from the same isolate, as first discovered by Wang et al. [24] (**Supplementary File S1**). SNP/kbp within a group ranged from 2.33 to 5.67 SNPs/Kbp, and between groups from6.67 to 9.07 SNPs/Kbp. Again, the exceptions were isolate AZ2, in which both haploid genomes belonged to Global A, and Pst130 in which low overall alignment to other genomes made SNPs/kbp comparisons unreliable. Surprisingly, this effect held true even with *Psh* isolate NDPSH, which carries a haploid Global A and haploid Global B genome (**Figures 1, 4**), rather than forming a *Psh*-specific outgroup, indicating that this split may predate the division of *Puccinia striiformis* into the present *formae speciales*, or else imply ongoing nuclear exchange between the groups.

**Figure 1.**
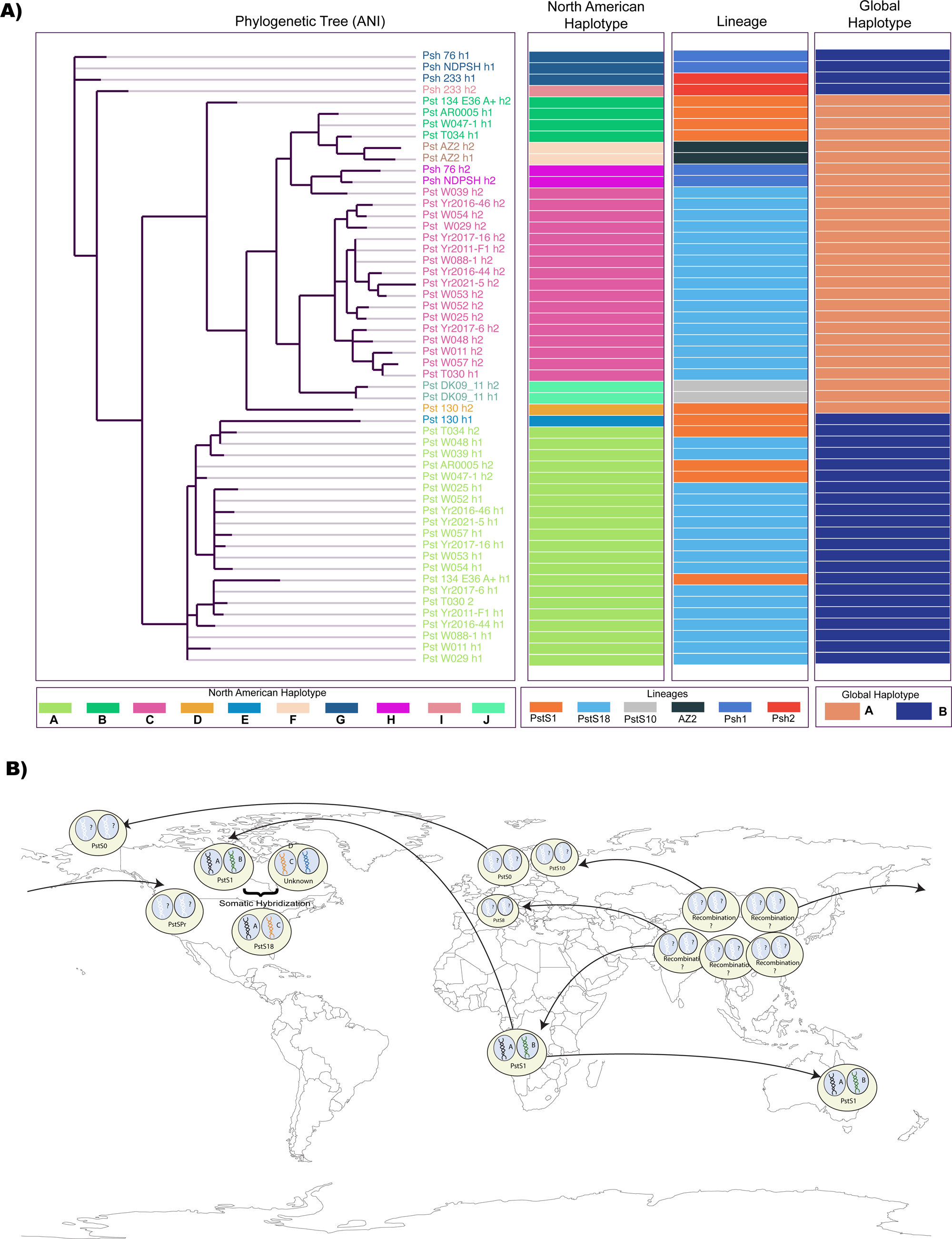
Relationships between North American *Puccinia striiformis* f. sp. *tritici* lineages. **A)** Phylogenetic tree of all phased haploid genomes in our study, based on Single Copy Core gene average nucleotide identity (ANI) as in Figure 4. Following the tree are coloured bars indicating three ways of classifying these genomes. From left to right: North American haplogroup, describing the ancestry of the clonal haploid genome as identified in this study. Lineage, the ancestry of the clonal isolate. Global haplogroup, which of the two conserved fundamental *P. striiformis* haplogroups the haploid genome falls into. **B)** Spread of selected *Pst* lineages around the world. PstS18 is descended from PstS1 via somatic hybridisation with an unknown lineage. The centre of diversity for *P. striiformis* is the Himalayan region, and genetic novelty is largely believed to originate there and spread outwards. Of note are PstS1, an old lineage which has spread across multiple continents and adapts independently to their conditions, and PstS7 and PstPr, lineages which recently emerged from the Himalayan centre of diversity and travelled to Europe and North America respectively.

**Figure 2.**
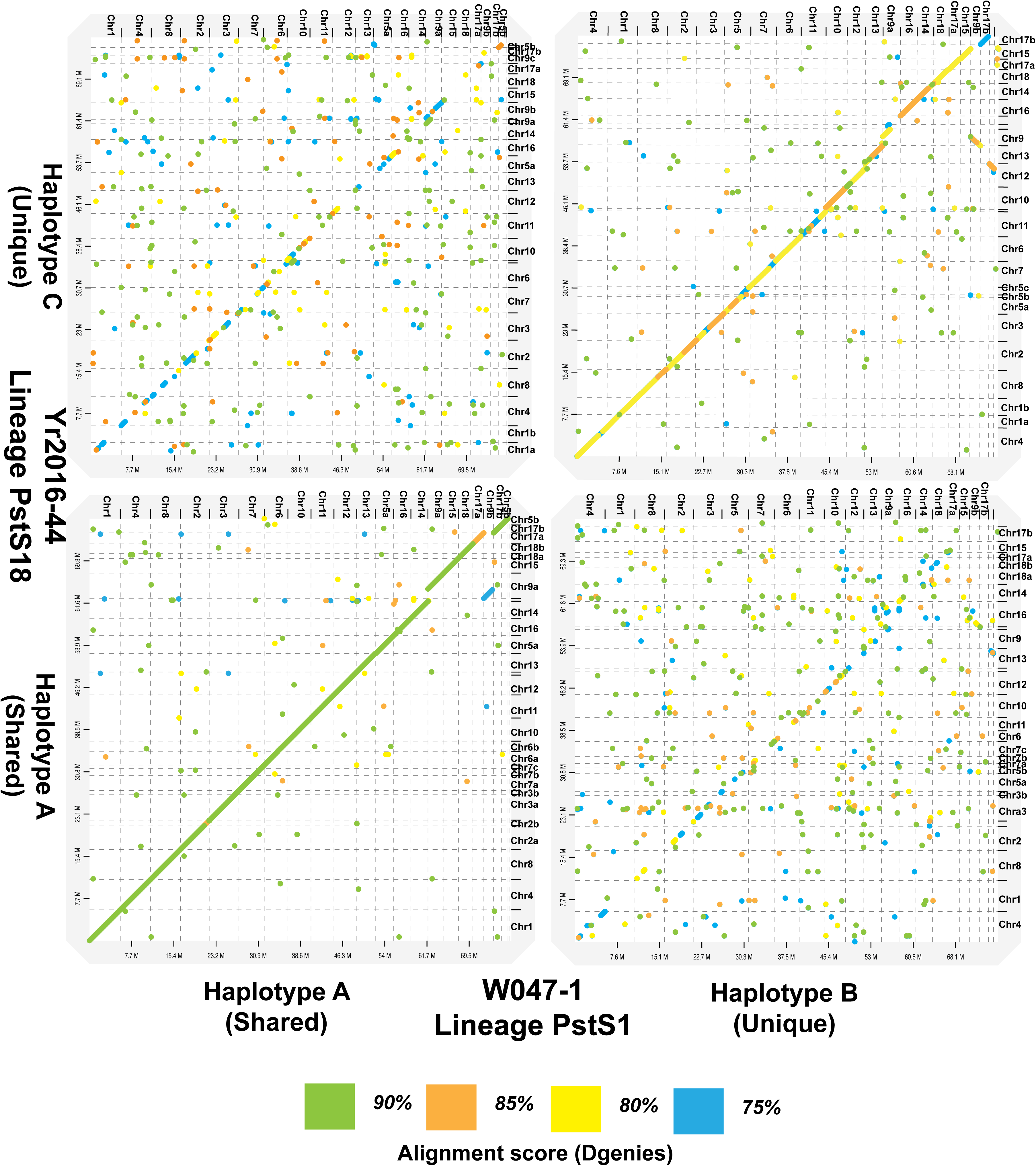
Concordance between phased haploid genomes of Yr-2016-44 (**A+C**), from lineage PstS1 and W047-1 (**A+B**), from lineage PstS18. The shared **A** haplotype is in near perfect agreement, whereas the **A-B**, **A-C**, and **B-C** haplotypes exhibit substantial sequence divergence. Figure generated using D-GENIES and visualised using Adobe Illustrator. Each of the 18 unscaffolded *Pst* chromosomes is captured in both assemblies to avoid overfitting the data to our hypothesis, with sub-sections of chromosomes described as a,b,c.

The most similar haplotypes were Yr2016-44 and W047-1, which are from lineages *PstS1* and *PstS18*, shown in this paper to be related by somatic hybridization. The second closest pair of genomes were W047-1 and the Australian isolate Pst134 E36 A+, both members of the *PstS1* group. (**Supplementary File S1**).

### Phased genome analysis reveals the origins of *PstS18* in a somatic hybridisation event

In a previous study [16], we found that lineages *PstS1* (isolate W047-1) and PstS18 (isolates Yr2016-44, T030) share a single mating-type. These lineages were previously believed to be a single lineage based on overall nuclear identity. Observing that their most similar haplotype exhibits 0.04 SNPs/kbp in our comparison, approximately 58x fewer than the next-most closely related haplotypes, while their complementary nuclear genomes exhibit a more usual 2.64 SNPs/Kbp, we directly compared their nuclear genomes and demonstrated that lineage *PstS18* (formerly *PstS1-related*) is descended from East African lineage PstS1 via somatic hybridisation with a third, unknown lineage (**Figures 1, 2**). Alignment of phased genomic contigs from isolates Yr2016-44 (PstS18) and W047-1 (PstS1) revealed that the shared haplotype (haplotype A) matches in its entirety, and exhibits no evidence of recombination, assortment, or rearrangement along or between chromosomes. The non-identical haplotypes (A-B, A-C, and B-C) exhibit between 120,000 and 375,000 SNPs, amounting to between 0.13% and 0.42% difference respectively (**Supplementary File S1**). The non-shared haplotypes B and C both fall into the same global haplotype lineage (Global A) but are as distinct from one another as they are from any other member of that group. No sample in our collection or in publicly available sequence data was a good match for haplotype C, and so the second parent of the somatic hybridization remains a mystery.

### Pseudo-phasing allows additional information to be gleaned from non-phased genomes

Given these results, and the established literature, we wanted to investigate how each distinct haploid genome (e.g., A, B, and C) has evolved across the lineages PstS1 and PstS18. After determining the lineages of our remaining *de novo* sequenced *Pst* genomes, we performed “pseudo-phasing” to enable further pan-genomic analyses utilising the distinct haploid genomes. Presuming that the isolates are the product of asexual reproduction within a lineage, contigs were aligned to the phased reference haplotypes from that lineage in order to produce pseudo-phased assemblies which can be compared as if they were phased using Hi-C data from that isolate. Critically, the resulting alignments were investigated for evidence of recombination or poor-agreement to the purported reference genome which would invalidate this approach, and this process revealed that pseudo-phased assemblies exhibited good overall coverage (>90% of the expected genome size) and SNP variation within haplotypes ranging from 384 SNPs to 9810 SNPs (0.042 SNPs/kbp to 0.1 SNPs/Kbp), an order of magnitude smaller than between the closest non-clonal haplotypes, validating our approach to differentiate haplotypes this way. After pseudo-phasing, contigs were scaffolded using RagTag and the Pst 134 E36 A+ genome. Manual pseudo-phasing was performed first to minimise the chance of contigs being placed in the wrong genome based on limited or arbitrary similarities, as RagTag is not intended for use as a phasing tool.

The pseudo-phasing approach was also taken with our two non-phased *Psh* samples. Psh-76, matched well with both haplotypes (G and H) of the phased NDPSH genome. However, H233 exhibited good matches to only a single haplotype (G) of the NDPSH genome. The remaining contigs were therefore assigned to a new alternate haplotype (I). As with *Pst*, the matching haplotype exhibits high alignment coverage and minimal SNP variation, leading us to conclude that these isolates are also related via an ancestral somatic hybridization event. Fascinatingly, isolate H233’s alternate haplotype I is in the same global group (Group B) as the primary haplotype G (**Figure 1A**) which, in the context of isolate AZ2, may also indicate a history of sexual recombination in North American *Psh* isolates.

In total, across our isolates and the publicly available phased reference genomes, we identify the following shared haplotypes which will be used throughout (**Table 1**).

A+B: W047-1, 3 additional North American *PstS1* isolates. 134E36 Australian reference Isolate.

A+C: Yr2016-44. 21 additional North American *PstS18* isolates (Including T030). D+E: Pst130

F: AZ2

G+H: NDPSH. 1 additional North American *Psh* isolate

G+I: H233

J: DK0911

In the global context [24], haplotypes B, C, D, E, F, H, and J are in global haplotype A, and haplotypes A, G, and I are in global haplotype B.

### Pan-Genomic analysis reveals distinct per-haplotype gene content of *P. striiformis* isolates

To compare gene family evolution across the North American pan-genome, we took the entire set of gene clusters (e.g. sets of genes which can be grouped by orthology or homology) present across all genomes and divided it into single-copy core (SCG), core, accessory, and singleton gene clusters. The SCG and core gene clusters are formed of the conserved set of gene families found in all haploid genomes as single or multicopy units respectively, while accessory genome consists of the dispensable component that is present in a subset of genomes and singleton genome is unique to only one haplotype out of all the analysed genomes. This approach can be scaled to distinguish between the features which present as core and accessory when comparing: all *P. striiformis* samples, *Pst* and *Psh* alone, a given *Pst* lineage, or a given *Pst* haplotype. It should be remembered that we compare 9 haplotypes from 7 lineages, with haplotypes unevenly distributed between lineages and so the degree to which a gene may be appear to be core or accessory will depend on the particular comparison being made. In particular, individuals with only a single representative such as DK0911 (PstS7/Warrior) will contain a high number of “unique” genes as per-lineage and per-individual uniqueness are combined.

Taking the full set of 27 phased and pseudo-phased isolates including previously published reference material, this gives 54 haploid genomes from 7 identified lineages (PstS1, PstS18, AZ2, Pst130 PstS7, PshS1, PshS2). The pan-genome is shown to be formed by a total of 16,145 gene clusters with 872,265 annotated genes across 18 scaffolded chromosomes (**Supplementary Figure S2**), that included 235 (1.5%) SCG, 2,271 (14.0%) Core, 10,763 (66.7%) accessory, and 2,876 (17.8%) singleton gene clusters (**Figure 3A**). A cladogram was generated by a phylogenomic analysis with 235 SCG gene clusters (**Figure 3B**) which agrees with our previously asserted relationships between individual haploid genomes: Haploid genomes are divided into two major ancestral groups, and that each haploid genome within an isolate is more related to other members of its ancestral group than to its complementary genome in that isolate. This approach also validated our previously manually assigned haploid lineages, designated A-I, as they were fully recaptured by the topology generated in this independent approach (**Figure 3B**).

**Figure 3.**
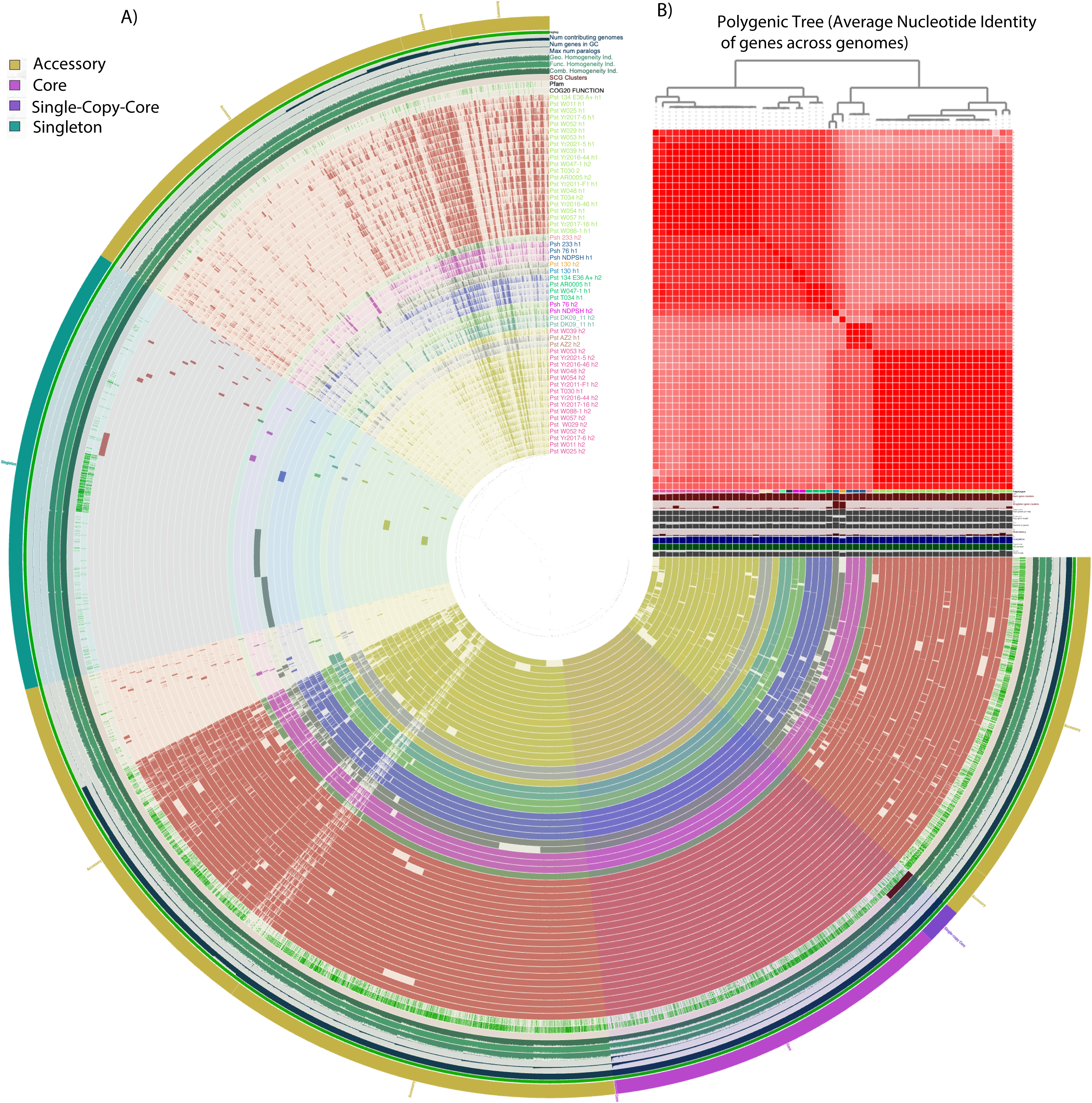
**A)** Pan-genome analysis. Clustering of 54 *Pst* and *Psh* haplotype genomes is based on the presence/absence patterns of 16,145 pan-genomic clusters. The genomes are organized in radial bins of the outside layer as single-copy core, core, singleton and accessory gene clusters (Euclidean distance; Ward linkage) which are defined by the gene tree in the centre. Genomes of haplotype A, B, C, D, E, F, G, H, I, J are arranged by phylogenetic tree of average nucleotide identity (ANI) and colored by red, blue, yellow, dark gray, light gray, purple, light green, yellow and cyan, respectively. Line segments indicate single gene clusters. The outermost circles carry the information regarding gene cluster located on chromosome (RagTag), the number of genomes containing specific gene cluster, number of genes in gene clusters, maximum number of paralogs, functional, geometric and combined homogeneity indexes, distribution of single-copy gene, Pfam and COG20 function. The layers below the ANI heatmap display from the bottom to top: total length of the genomes, GC content, genome completion, redundancy of each genome, number of genes per kbp, number of singleton gene clusters, total number of gene clusters and haplotypes. **B)** Heatmap denotes correlation between the genomes based on average nucleotide identity values calculated using pyANI.

The accessory component forming more than half of the pan-genome indicated that the set of genes conserved across all nuclei is much lower than had previously been thought. However, a proportion of genes marked as accessory were only absent in one or two unrelated genomes which might be attributable to miss-assembly and inflate this number somewhat. Additionally, any amount of presence-absence variation within a lineage or between the two represented *forma speciales* is enough in this model for a gene cluster to be categorised as accessory, and it is likely many of these genes fulfill equivalent functions. Across each haploid lineage of A, B, C in *Pst* and G, H, I in *Psh*, the number of present accessory gene clusters varied, but statistically each lineage exhibited a consistent 8,000-9,000 accessory gene clusters present while haplotypes of D, E, F, J and K harbor over 10,000 accessory gene clusters on average (**Figure 4B**), which concurs with the explanation that low sample-number inflates uniqueness. The number of total genes in any given *P. striiformis* genome appears to be stable at around 15,000-16,000, with around 60% of these genes being either dispensable or interchangeable.

**Figure 4.**
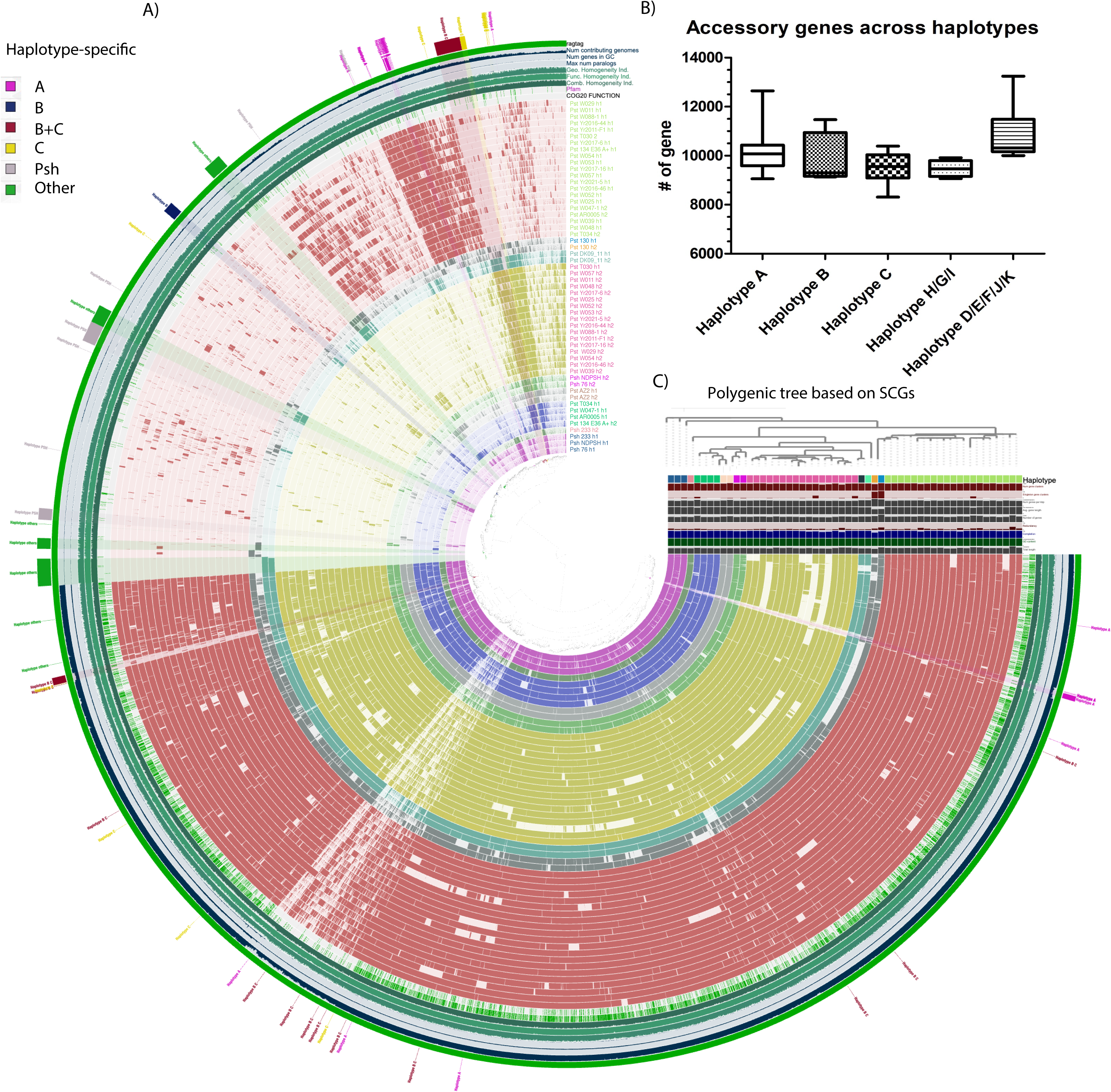
**A)** Split pan-genome analysis. Clustering of genomes based on the presence/absence patterns of 10,763 accessory gene clusters. The genomes are organized in order of gene cluster frequency and highlighted by radial bars as haplotype-unique gene clusters [Euclidean distance; Ward linkage] that are defined by the gene tree in the centre. **B)** The boxplot denotes the number of accessory genes in single haplotype - A, B, C, and combined *Psh* and *Pst* haplotypes - H+G+I and D+E+F+J. **C)** The haplotypes are colored based on the pan-genome tree classified by 235 single-copy core gene clusters.

To evaluate the genomic basis of lineage for signs of which gene families drive lineage specificity and evolution the accessory gene cluster was evaluated for differences between haplotypes A, B, C, G, H, and I, representing the circulating North American *P. striiformis* population. 6,535 (40.5%) of accessory gene clusters were found in over 80% of all haploid genomes with no clear pattern underlying presence/absence (**Figure 4A**) and could perhaps be regarded as an extended core genome. 3,425 (23.9%) gene clusters display a presence/absence pattern which is haplotype-specific, between one or more of haplotype A (PstS1 and PstS18), haplotype B (PstS1), haplotype C (PstS18), haplotypes G+ H/I (*Psh*). The genes which are unique and always conserved within a haplotype define the haplotype and haplotypes B and C (e.g., *Pst* Global A) exhibit conserved genomic differences compared to the *Pst* Global B (haplotype in PstS1 and PstS18).

The largest category of gene clusters is the set which are not fully conserved but are uniquely present in a given haploid lineage (**Figure 4A, Supplementary File S4)**. This gene set may explain why clonal lineages present so much phenotypic variation: there are a high number of unique genes which are demonstrably plastic inside a lineage. Noticeably, the largest set of such genes belongs to the A haplotype, shared between *PstS1* and *PstS18*, meaning that genome reorganization following the somatic hybridisation event leading to the emergence of *PstS18* is still ongoing in the clonal population.

### Core gene families regulate essential biological functions while nearly half the pan-genome is predicted to be dispensable

To gain more insights into how gene function and dispensability/uniqueness are connected, the core, accessory, and singleton gene clusters were further annotated using COG and Pfam classes (**Supplementary file S4, S5 and S6**). Relative to overall genome content, the 2,506 core gene clusters are enriched in the following categories (**Supplementary file S5**): carbohydrate transport and metabolism (12.8%), translation, ribosomal structure and biogenesis (12.4%), amino acid transport and metabolism (9.1%), posttranslational modification, protein turnover and chaperones (8.9%), replication, recombination and repair (8.4%), lipid transport and metabolism (8.3%), and signal transduction mechanisms (7.6%). The classes for coenzyme transport and metabolism (5.9%), inorganic ion transport and metabolism (5.0%), cell wall/membrane/envelope biogenesis (4.5%), energy production and conversion (4.4%), transcription (4.3%), secondary metabolites biosynthesis (3.9%), and transport and catabolism (3.3%) are also moderately abundant in the core. As rusts are biotrophic fungal pathogens, their core genomes are thus enriched in genes for sensing external stimuli, efficient uptake of nutrients from host and cellular signalling and growth.

Perhaps unsurprisingly, many of the singleton gene clusters did not have a predicted function, indicating that the gene models may be incomplete, or that their function has not been established due to their overall rarity as a class of protein-encoding gene. (**Supplementary File S6**).

The accessory genome comprises the majority of the pan-genome’s predicted coding genes and was not enriched for any particular gene categories; however, when broken down into per-haplotype analyses interesting patterns are revealed.

### Haplotype-specific functional profiles might explain differences in fungal virulence and host adaptation

Following on from this, we wanted to investigate the relationship between specific accessory gene clusters and genes and individual lineages in order to understand what drove the evolution of PstS18 (A+C) in North America to be distinct from, and more successful than PstS1 (A+B). The number of unique gene clusters/genes identified per haplotype or group were A (57/2150), B (47/369), C (32/1111), B plus C (156/6314), *Psh* isolates G, H, I (145/530), and others (e.g. reference genomes DK0911, Pst130 and AZ2) (D, E, F, J) (331/1738) (**Figure 4A, Supplementary File S5**). We defined uniqueness as those gene clusters/genes are found in over 80% of that specific haplotype or group and found in at most one genome outside that haplotype, being present in nearly all examples and absent outside of it. We further substantiate the functional roles of haplotypic-specific gene clusters from pan-genome by cataloging over 30 gene clusters associated with one or more haplotypes from A to J, annotated with Pfam and COG domains and linked to fungal virulence, stress response, and host adaptation (**Table 2 and Supplementary table S7**). Six specific gene clusters in haplotype A (**Table 2**) were identified as encoding proteins with predicted functions including DDE superfamily endonuclease, heterokaryon incompatibility protein (Het-C), Myb/SANT-like DNA-binding domain, reverse transcriptase, and transcription factor AFT and AT hook motif. These domains are functionally linked to post-translational modification, genome stability, and transcriptional regulation which could participate in essential pathways for fungal survival and pathogenicity. In haplotype A plus *Psh* haplotypes, domains of RecD & YpbB and Dut involved in genome maintenance and transcriptional control ensure chromosomal stability and regulate gene expression under host-induced stress. Interestingly, two gene clusters, GC_00011751 and GC_00010102, were associated with haplotype A and G and identified to encode proteins containing a conserved RING finger domain (Pfam: PF13639) of E3 ubiquitin ligases, implicating a functional role in the modulation of host immune signaling pathways (**Table 2**).

**Table 2.** Functional roles of haplotype-associated Pfam domains and COG20-group proteins in fungal virulence and evolution. The gene clusters enriched in different haplotypes were generated by frequency table of functional enrichment test implemented by Amy Willis in R (https://github.com/adw96), and filtered by using a Generalized Linear Model with the logit linkage to set a threshold of enrichment score > 20 and p/q-value < 0.05 for each function truly enriched in certain groups.

Haplotypes B, C, E and H (global haplotype A) shared six unique clusters, which included functional domains of dynactin p62 family, Alanine-zipper lipoprotein, NuoL, PRP8 domain IV core, alanine-zipper lipoprotein and AspB with predicted functions in vesicle trafficking, membrane structure maintenance, respirational proton transport, and RNA splicing and amino acid biosynthesis (**Table 2**). In haplotype C, unique gene clusters found in over 80% of haplotype C genomes were N-terminal SNF2-related domain and RNase H-like domain in reverse transcriptase (**Table 2**). In fungal pathogens, SNF2-like proteins acting as part of a chromatin remodeling complex are often linked to the regulation of effector genes and stress-responsive pathways [25,26]. This domain might allow *Pst* to quickly adjust its gene expression in response to host defenses, contributing to its ability to expend the host range. Moreover, RNase H-like domain could facilitate genome plasticity by promoting retrotransposition in a reverse transcriptase framework, allowing the fungus to generate genetic diversity possibly contributing to host or environmental adaptions. It may also play a role in silencing host defense genes or regulating fungal effectors through RNA interference-like mechanisms [27]. Among the *Psh* isolates (haplotypes G, H, I), three unique virulence-associated gene clusters were identified which always distinguished them from *Pst* (**Table 2**). These genes encode functional domains of zinc knuckle domain, RNase H and Tet-like 2OG-Fe(II) oxygenase superfamily known to modulate effector gene expression via epigenetic regulation and RNA binding in the pathogen and the host [27,28].

Besides unique gene clusters present in single, double or triple haplotypes (**Table 2**), more gene clusters associated with more than 3 haplotypes have been identified and their functional roles in pathogenicity and pathogen-host interaction have been explored (**Supplementary table S7**). Several domains are directly linked to pathogenicity, such as the AT hook motif and SWIM zinc finger, which facilitate effector targeting to host nuclei and regulate stress-responsive transcription [29,30]. Structural components like the Dynactin p62, Erg28-like protein and Alanine-zipper lipoproteins contribute to cytoskeletal transport, membrane integrity and host interaction [31], while PMT C-terminal TMM modifies surface glycoproteins to evade immune detection [32]. Domains involved in genome maintenance and transcriptional control such as Mif2_N, PIF1-like helicase, RecQ helicase, and SNF2 helicase ensure chromosomal stability and regulate effector gene expression under host-induced stress [33]. Metabolic and stress-response domains like CarB, AspB, and glutamate 5-kinase support fungal proliferation and osmotic tolerance within host tissue. Virulence signaling can occur through Cdc37 and Spt20 SEP domain, which stabilize kinases and remodel chromatin to activate infection-related pathways [35]. RNA processing domains like PRP8 and RNase H/H-like facilitate transcript maturation and genome adaptability through retro-transposition [28]. Additionally, ZntA and CopZ mediate metal ion detoxification, allowing rust fungi to withstand host-imposed stress [36]. Notably, domains such as DUF1588, DUF6589 and Tet-like 2OG-Fe(II) oxygenase are enriched in pathogenic fungi and may represent novel virulence factors or epigenetic regulators of effector expression. Interestingly, two gene clusters of aspartyl protease associated with haplotype groups of A, F, J and K or C with D, E, F, J, K (PST130, AZ2 and DK0911) have been characterized as a secreted fungal protein to facilitate the transformation of yeast cells into hyphae and engage in biofilm formation, invasion and degradation of host cells and proteins [37]. In *Arabidopsis thaliana*, aspartyl protease can couple with molecular co-chaperones to trigger autophagy and plant defense [38]. Recent research suggests aspartyl proteases and their isoenzymes could be a crucial target to disrupt fungal pathogenesis such as adhesion, hyphal development, host invasion and immune evasion [39,40].

Notably, *Pst/Psh* isolate-specific gene clusters and genes i.e., those uniquely present in a single haploid genome, were prominently featured in the split pan-genome analysis comprising 2,876 singleton gene clusters (**Figure 5A)** and found to evenly distribute across all the chromosomes rather than being present in syntenic clusters on propsed pathogenicity islands (**Figure 5B, Supplementary File S2**). Across 10 distinct isolates, Pst130.1 (D haplotype) and Pst130.2 (E haplotype) exhibited the highest number of gene clusters/genes as 614/626 and 561/572, corresponding with their status as an outgroup from a very divergent lineage compared to other samples, as well as the fragmented nature of those genomes. Pst134E36.1 and Pst134E36.2 demonstrated consistent profiles, each with 174/174 and 132/132 unique gene clusters/genes, placing them as evenly divergent from the other PstS1 lineage samples in our analysis as expected from Australian isolates adapted to that environment. Interestingly, two genomes of C haplotype PstT030.1 and PstW052b yielded 197/197 and 233/233 unique gene clusters/genes while their corresponding genomes of the A haplotype: PstT030.2 and PSTW052a showed very low levels of unique gene clusters/genes as 7/7 and 7/8, respectively. Thus, the abundance and distribution of these isolate-specific genes can differ across different haplotypes within the same isolate, and again the PstS18 lineages exhibits a great deal of variation implying ongoing nuclear reorganisation across North America as the lineage continues to adapt. We speculated that the origin of these additonal genes may be from recombination or horizontal gene transfer from another isolate, but were unable to find any matches in our collection to indicate this, and the genes were not in a single location as would be expected from a donation of a single chomosomal region such as via recombination or transfer.

**Figure 5.**
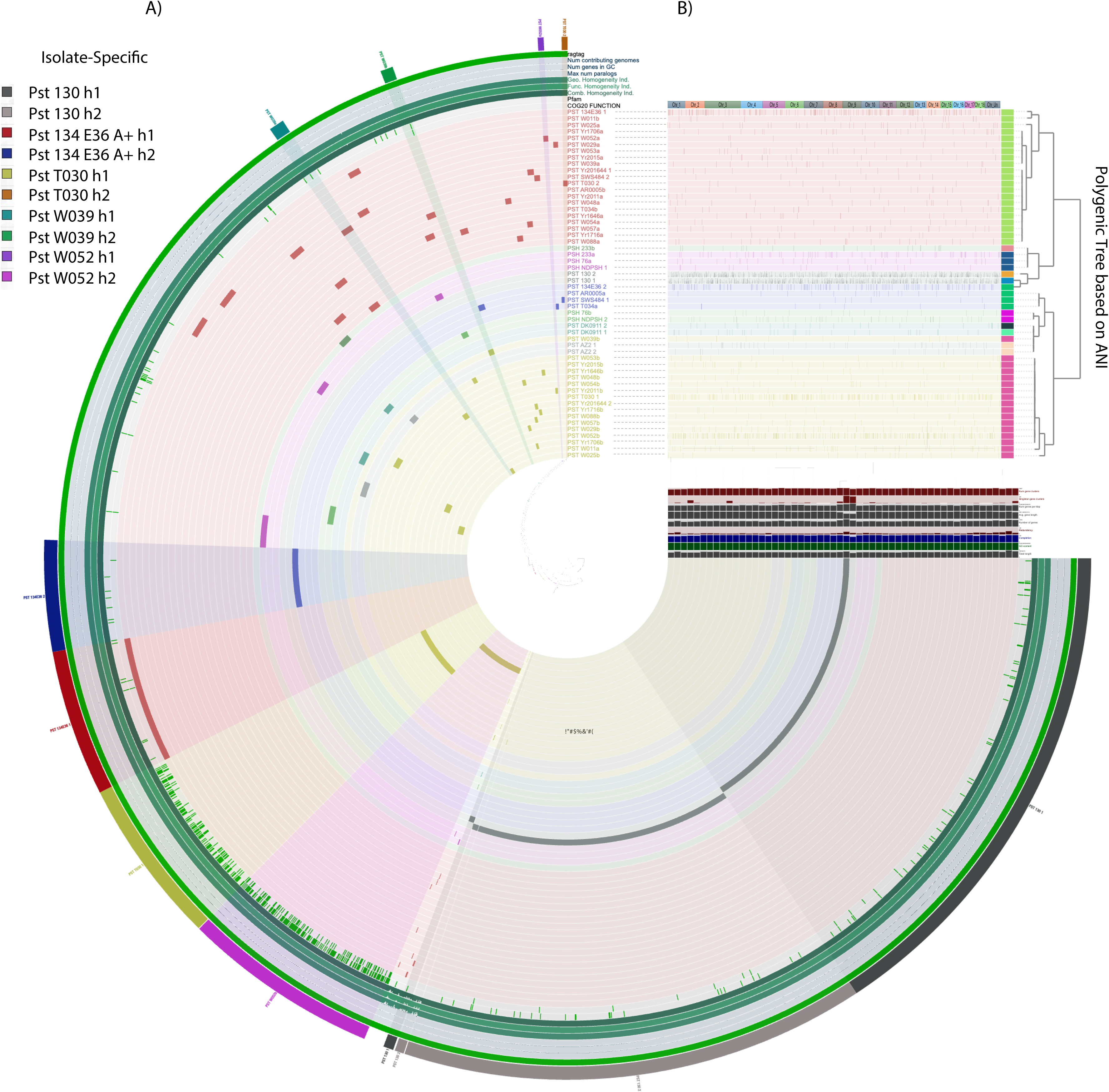
**A)** Split pan-genome analysis. Clustering of genomes based on the presence/absence patterns of 2876 singleton gene clusters. The genomes are organized in order of gene cluster frequency and highlighted by radial bars as PST isolate-unique gene clusters [Euclidean distance; Ward linkage] that are defined by the gene tree in the centre. **B)** The phylogram layer data denotes the presence/absence of singleton genes along 18 PST/PSH chromosomes (Chr_1 to Chr_18) or unsorted locations (Chr_Un). Genomes of haplotype layers are arranged by phylogenetic tree of average nucleotide identity (ANI).

### High variability of North American stripe rust field populations has been shaped by diversity in the pan-genome

All isolates taken from North American fields in our study were phenotyped using our in-house differential panel of 18 single R-gene lines (**Table 3**). Isolates were found to belong to one of six races on this differential set, with race largely uncorrelated with lineage. Additionally, isolates belonging to PstS18 (especially T030 and W039) were observed to be highly aggressive, producing large numbers of spores on susceptible plants and having the widest ranging virulence. The observation that the North American *Puccinia striiformis* pan-genome exhibits high inter- and intra-lineage variability may go some way towards explaining the historically common observation that pathogen race and genetic lineage are not necessarily correlated – circulating clonal populations frequently adapt to overcome resistance genes or gain tolerance to fungicides.

**Table 3.** Isolates used in this study, and their origin, race, and virulence on differential wheat accessions. Some isolates are displayed with a WXXX (official) and additional (temporary) name.

## Discussion

We have constructed a North American pan-genome of *Puccinia striiformis*; the stripe rust fungus, focussing on the population of North America which is geographically and reproductively isolated from the other major circulating populations worldwide. Our analyses revealed that the predominant North American lineage PstS18 has its origin in a somatic hybridization event between PstS1 and an unknown third lineage, with the A haplotype being shared between them. We also show strong likelihood of somatic hybridisation in the related barley-infecting *forma speciales P. striiformis* f. sp. *hordei*. Together with other results from recent years [19–24] this data suggests that somatic hybridisation may be a crucial driver of rust pathogen evolution in the absence of a sexual host, permitting genetic novelty to arise and helping to balance for the expected loss of fitness or adaptability postulated to affect asexual species via Muller’s ratchet [41]. While somatic hybridization may be an uncommon event on a per-individual level, bearing in mind the billions of opportunities for hybridization in the field each growing season it may well be that somatic hybridization occurs regularly and simply does not often produce an offspring with an increase in fitness relative to either parent which therefore makes it extremely difficult to measure true hybridization using a limited and random sampling of isolates.

We determined that approximately 63% of the total gene content in the pan-genome, and 40% of genes on a per-isolate basis exhibited presence-absence variation. This is similar to previous reports of up to 25% difference between the two genomes of a single isolate [16]. Differences between the two haploid genomes of the same isolate were high in all cases except for the recent sexual recombinant AZ2 and the *Psh* isolate H233, which raises the intriguing question of if this genomic signature supports recent sexual recombination in these isolates too. Coupled with earlier findings that haploid rust genomes worldwide fall into one of two global groups (Group A and B), these data overall suggest that the two rust nuclei may carry distinct nuclear complements and be evolving to fill separate roles in the pathogen’s biology [24]. We also identified pervasive presence-absence variation in gene content even within a single haplotype in a lineage. This suggests that asexually reproducing rust populations still maintain a pool of genetic diversity which can account for the continual emergence of new races in the field. Particularly isolates T030 and W052 of lineage PstS18 feature >400 unique gene clusters between them, which raises additional questions as to the origins of genetic novelty in clonal rust lineages in the absence of an apparent recombination event. These differences, however, are minor in comparison to the degree of unique gene family presence/absence variation between haplotypes (**Figure 4**).

We identified consistent differences between isolates which have historically been grouped together, particularly members of the PstS1 and PstS18 lineages, which we attribute to the resolution afforded by our approach. In the genomics era, other methods of categorisation include amplicon sequencing of barcodes such as ITS or similar single markers, sets of genomic markers ranging from KASP / RFLP / SNP sequencing and the MARPLE approach which is a form of enriched / targeted genome sequencing, and WGS / RNAseq combined with variant calling and phylogeny construction. In each case, the approach is limited by the availability of markers and in particular the inability to distinguish between the two haploid genomes without extensive foreknowledge of their content. Indeed, for many years it was assumed that PstS18 was a subset of PstS1 based on their overall genomic similarity. With our phased assemblies we can account for this as being due to 50% of the diploid genome being identical, and the remaining 50% radically different. Approaches that collapse diploid genomes together can err on the side of treating heterozygous variation as equivalent to both constituent pieces. Without phasing, and in the circumstance of a freely intermixing, sexually recombining diploid species this may be more reasonable, but in this case, it prevents a proper understanding of the species. In this pan-genome, we demonstrate that the differences between lineages of *Pst* can be attributed to conserved differences in individual haploid genomes that are consistently maintained across a population, and that only separating out these genomes allows one to properly investigate gene flow and genome organisation within the species. PstS1 isolates from North America and Australia show marked differences despite both originating from the same population in east Africa [15]. Whether this represents a subset of diversity from that founder population or genetic novelty arising in each continent will be a fascinating question to resolve as additional genomes from across the world are sequenced in this way.

Analysis of the distribution of genes across fungal genomes by haplotype and lineage found each haplotype could be defined by between 300-3,000 unique genes conserved within that haplotype (**Figure 4B**). Expanding the criteria to include genes conserved across related haplotypes produced different sets of genes, yet in similar orders of magnitude. We used a criterion of >80% conservation to describe conservation within a haplotype, but alternative cutoff points or considering multiple related haplotypes may reveal different evolutionary signatures based on the biology of the pathogen. Notably, within haplotype A, shared by isolates in lineage PstS1 and PstS18, genes were identified that are additionally shared across all isolates, shared only by PstS18 isolates, and that separate North American PstS1 and PstS18 from the Australian PstS1 sample 134E36. Further phased genome sequencing of PstS1 isolates from around the world will be needed to properly understand ongoing change and genomic diversity within this lineage and its connection to fitness for the local environment. The origin of PstS18 in a somatic hybridization event offers a clear example of an A/B test (in this case, a “B/C” test) whereby the total replacement of one nuclear genome for another provides a fitness advantage which can be directly linked to the gene content of that genome.

The sequencing of a pathogen genome represents a landmark development in our ability to make informed decisions about avenues of research (e.g., into fungicide tolerance and disease resistance), as well as to understand the evolutionary history of the species and how that relates to its effects on agriculture. The origin, behaviour and movement of these species worldwide is of primary concern to growers and scientists alike, as significant time and money expenditures are made for disease prevention measures such as variety control, border biosecurity, and field monitoring for novel disease strains. This first *Puccinia striiformis* pan-genome will drive a major leap forward in our ability to understand the species as a whole. As accurate long-read genome sequencing and conformation capture sequencing becomes more and more accessible, the ability to incorporate fully phased and sequenced genomes from around the world will only increase, and it will be critical to examine what patterns have affected the genetics of the *P. striiformis* populations in different locales. Crucially, are signs of somatic hybridization evident in many areas, or primarily in North America? Are the core and dispensable genome components conserved across the entire species, or do some regions exhibit very different patterns in this regard? And are the two worldwide haploid genome groups truly distinct, and each providing fundamentally different genetic material to the pathogen, or is this apparent division simply a result of sequencing bias from countries with large single-lineage populations?

## Materials and Methods

### Fungal isolates and purification

A total of 25 *Puccinia striiformis* f. sp. *tritici* (*Pst*) and 3 *P. striiformis* f. sp. *hordei* (*Psh*) isolates have been newly characterized during this research (**Table 1**). All these isolates were originally obtained from naturally infected commercial wheat and barley crops during annual disease surveys. Purification of the isolates was done by transferring urediniospores from a single pustule or single stripe to a leaf of the cultivar Avocet grown to the two leaves stage (12-14 days old) for *Pst* and to the cultivar Topper with the same age for *Psh*, using a fine tip needle. Avocet and Topper seeds were grown in 10-cm diameter plastic pots filled with potting mix. All inoculated plants were kept in dew chamber at 11°C for 24 hours in the dark and then transferred to Conviron PGR-15 Reach-In growth chamber (Controlled Environment Limited, Winnipeg, Manitoba, Canada) with 16 hours photoperiod (16°C day/12°C night). For increasing urediniospores of pure isolates obtained from above procedure, 14 days old Avocet and Topper plants were inoculated with 20 mg of pure urediniospores suspended in 5-6 mL NOVEC 7100 engineered fluid using an airbrush compressor system with 0.3 mm Nozzle. Inoculated plants were kept in the dark inside a dew chamber at 11°C for 24 hours and then transferred to clear plastic sterilite latch box in a Conviron growth chamber (with a 16 hour photoperiod 16°C during day/12°C during night) to prevent cross contamination among isolates.

### Urediniospore harvesting and storage

After 12–14 days, the emerging urediniospores are mature and were harvested over several days. To harvest spores infected leaves were shaken over a fresh sheet of aluminium foil, then then transferred to 2 mL low binding Eppendorf microcentrifuge tubes by creasing and tipping the foil. Spores were dried in a desiccator filled with silica gel desiccants for 5-7 days with daily shaking of the Eppendorf to ensure even coverage and prevent mould. For short term storage spores were stored in a refrigerator at 4°C. For long term storage Eppendorf tubes containing spores were stored in a deep freezer (−80°C).

### Virulence phenotyping of *Pst* isolates

Virulence phenotypes of the *Pst* isolates were obtained on a set of 18 Yr single-gene differentials for commonly used resistance genes *Yr1*, *Yr5*, *Yr6*, *Yr7*, *Yr8*, *Yr9*, *Yr10*, *Yr15*, *Yr17*, *Yr24*, *Yr27*, *Yr32*, *Yr43*, *Yr44*, *YrSP*, *YrTr1*, *YrExp2*, *Yr76* (*YrTye*) and *YrA* and Avocet susceptible (*AvS*). Two weeks old seedlings were inoculated with urediniospores as described above. Infection types (IT) were recorded 15–22 days after inoculation and scored on a 0–9 scale [42]. The IT data on both first and second seedling leaves were recorded. An isolate was considered avirulent on a specific differential line when the plants show IT 0-6 or virulent when showing ITs 7-9. In case of an isolate producing different ITs on the first and second leaves, the IT of the second leaf, instead of the first leaf, were used for designating the isolate to a race. Races were named based on Octal code and Virulence formula on *Yr* genes [42].

### HMW DNA extraction and sequencing

High molecular weight (HMW) genomic DNA was isolated for PacBio sequencing using our previously published protocol [43]. The integrity and size-distribution of DNA was determined on TapeStation (Agilent, Canada) (**Supplementary Figure S3**). DNA concentration was measured using a Qubit™ 4 Fluorometer (Thermo Scientific™, Canada) and adjusted to 100 ng/µL using TE buffer. To optimize fragment size for meeting HiFi requirements, purified HWM DNA was sheared using g-TUBE (Covaris, Massachusetts, USA) and size selected to target fragments between 10 kbp–30 kbp by BluePippin (Sage Science, Massachusetts, USA) using manufacturer recommended protocols. Sequencing libraries were prepared using the SMRTbell Prep Kit 3.0, and barcoded overhang adapter kits 8A and 8B (Pacific Biosciences, Menlo Park, USA) following the manufacturer’s protocol (102-166-600 REV02 MAR2023) and then sequenced on 8M SMRTCells on a Sequel IIe Instrument (Pacific Biosciences). Each SMRT Cell yielded 20-35 Gbp of HiFi data for 5 rust isolates and the depth of sequencing expected to be 50X coverage of an individual *Pst* or *Psh* genome. Finally, the sequencing reads were then corrected to the FASTQ format and delivered by the sequencing platform. Sequencing was conducted using a PacBio Sequel II, IIe using Binding kit 3.2 according to the manufacturer’s instructions. Finally, the sequencing reads were then corrected to the FASTQ format and delivered by the sequencing platform.

### Hi-C preparation and sequencing

Fresh urediniospores were treated with formaldehyde and prepared for Hi-C library preparation according to the following protocol: ∼500 mg spore samples previously stored at −80°C were thawed and transferred to a 15 ml conical tube and left in a drawer overnight with lid ajar to rehydrate. Urediniospores were mixed with 1 ml/100 mg of 1% formaldehyde solution (5 ml for 500 mg) and vortexed for 20 mins. Glycine was added to a final concentration of 125 mM (46 mg for 5 ml) and the mixture vortexed for 15 mins, then centrifuged at 4,000 rpm for 10 mins to pellet the spores. Supernatant was discarded and the pellet was washed with 25 ml water, and then the wash step was repeated. The spore pellet was placed in a pre-chilled mortar and pestle with liquid nitrogen and ground to fine powder. The powder could then be transferred to a screw cap tube and stored at −80 °C before shipping on dry ice to Phase Genomics, USA, for Hi-C sequencing.

### Genome assembly

Raw PacBio sequencing data was evaluated for quality using FastQC v0.11.9 [44] and kat v2.4.1 [45]. Genomes were assembled using Hifiasm v0.19.5-r587 [46] with the additional options -s 0.45 --hg-size 85m. The isolates with H-iC data (W047-1, Yr2016-44, T030, W056, and NDPSH) were assembled using the additional -h1 and h2 options. Assembled genomes were subjected to an additional filtering step during gene prediction with Funannotate of removing contigs shorter than <30 kbp or with GC content over 70% or under 30%.

Additional QC on assembled genomes was performed by aligning them to the reference phased genome Pst 104 E36 A+ with mummer to validate that each chromosome has appropriate copy number, and visualising the distributions of *K*-mers in the raw and assembled reads with Kat to validate that the heterozygosity of each genome as proportional to the input data.

### Pseudo-phasing and evaluation of lineage

With the presumption that members of a lineage are clonal and should not exhibit recombination or independent assortment of chromosomes, we pseudo-phased the genomes without Hi-C data available by aligning the assembled contigs to the closest fully phased diploid genome and extracting contigs with a high degree of contiguity. Initial alignments were performed using minimap2 [47] in -asm5 mode, with samtools view, sort, and fasta used to re-extract contigs with a match to each genome. Samtools view was used to convert reads to bam format and drop unaligned reads, and once again to parse out reads matching one haploid genome using a bedtools index generated from that haploid genome. Newly pseudo-phased genomes were then realigned to each phased *Pst* haploid genome using the same parameters, and SNPs were called and evaluated using bcftools mpileup, call, filter, and stats. bcftools call used the parameters –ploidy 1 -m -v to identify polymorphisms in a haploid genome. Quality controls oriented around read depth and quality are inappropriate in the case of an already-assembled genome and so were not used for this step.

In cases where this approach produced pseudo-phased genomes with an apparently high number of SNPs relative to their reference, another alignment was performed using the MUMmer suite program nucmer [48] in -mum (maximal unique match) mode, and the resulting delta file filtered using delta-filter. Outputs were interrogated using show-coords and mummerplot to evaluate contigs for binning and to verify that the whole genome was reasonably well covered (e.g., each chromosome had a single matching contig for the majority of its length). Contigs with a minimum alignment agreement of 99% and alignment length of 1 Mbp were binned into the same haploid sub-genome as their closest match. Contigs without a clear single match were left unplaced. This approach was taken for T030, W054, W039, and H233. In the case of T030, W054, and W039 a high degree of agreement was found to the A haplotype, but manual re-alignment was needed to produce satisfactory matches to the C haplotype. In the case of H233, alignment was found only to the G haplotype, and other contigs were assigned to the I haplotype.

In three cases, the empirically determined lineage and phenotype (eg by sequence similarity and mating type, and by virulence phenotyping) did not match our expectations. These were isolates W047/SWS484, W088, and DK03-33. In these cases, the purified isolate in our collection used in the study was different to the sample as originally recorded. These isolates were re-named W047-1, W088-1, and DK03 to recognise that the isolate phenotyped and sequenced are not the original, but that they have behaved consistently since purification as a single isolate.

### Genome scaffolding

RagTag [49] was used to scaffold phased annotated (eg post-Funannotate) genomes to the closest available fully phased reference genome. In this way, Lineage PstS1, Yr2016-44, and T030 were scaffolded to the Australian isolate Pst 134 E36 A+, and members of lineage PstS18 to Yr2016-44. Scaffolds were joined together using a pre-defined repeat group of 100 N nucleotides. This approach was also used to scaffold the GFF3 files in order to compare synteny across predicted genes.

### Genome annotation

Genomes were annotated using the Funannotate pipeline [50].Phased and pseudo-phased genomes) were treated as individual genomes for the purpose of this process, but unphased genomes were grouped together. In brief, genomes were first sorted and filtered using Funannotate clean and Funannotate sort to remove small (<30 kbp,), unusual GC (>70%/<30%), and redundant (>95% identity) contigs from each assembly and output renamed contigs sorted by size. These genomes were then annotated for repeats using panEDTA [50], guided with CDS transcripts generated by running Funannotate predict on the unmasked W047-1 assembly in RNA-guided mode as below.

panEDTA outputs were used to softmask each genome with the provided EDTA make_masked.pl script, the softmasked genomes were then provided to Funannotate train, Funannotate predict, and Funannotate update along with RNAseq reads previously obtained from urediniospores of isolates BC-SP1, BC-SP2, W088-1, and USA_150.5333 [16,52].

After gene prediction and annotation, functional prediction was performed using local AntiSMASH v5.1.2 and InterProScan v5.66-98.0 installs, then combined with Pfam, UniProt, EggNog, COG, MEROPS, dbCAN, BUSCO, and SignalP using Funannotate annotate.

Gene annotations across all annotated assemblies were compared for basic statistics using Funannotate compare and orthofinder.

### Pan-genome analyses in Anvi’o

The pan-gene clusters were identified using pangenomics workflow in Anvi’o [53] and the genomes were organized based on the distribution of gene clusters using MCL algorithm into core, dispensable and strain-specific contents (Distance: Euclidean; Linkage: Ward). This method was also applied to determine singletons and single-copy gene clusters (SCGs). Singletons are defined as genes present only in single genomes. SCGs are represented by only one copy in each genome. The genes corresponding to amino acid sequences were annotated by BLASTp through Funannotate against the NCBI COG and Pfam database. Heatmap based on the annotated COG functions of the accessory and singleton gene clusters were then plotted in R. Data was prepared for Anvi’o from the outputs of Funannotate using custom python scripts to merge GFF3 files describing gene location with the predicted protein sequence of those genes in the appropriate input format.

## Data availability

Raw data and published genomes are available at NCBI under BioProject PRJNA1347092 and linked with BioSamples SAMN52837295 to SAMN52837321

## Acknowledgements

We are grateful to Mogens Hovmøller and Ben Schwessinger for their input, advice, and scientific discussions which added value to this project. The senior author (Gurcharn S. Brar) wholeheartedly remembers and dedicates this work and manuscript to his secondary school teacher, S. Chamkaur Singh Brar – a devoted chemistry lecturer and school principal (Govt, Model Secondary School, Bagha Purana, Moga) who had an enduring impact on his character and work ethic.

## Author contributions

SH: Samuel Holden. ML: Meng Li. MA Mehrdad Abbasi. RB: Ramandeep Bamrah. SHK: Sang Hu Kim. SF: Sean Formby. JB: Janice Bamforth. SW: Sean Walkowiak. AF: Andrew Friskop. USG: Upinder S. Gill. BM: Brent McCallum. HSR: Harpinder S. Randhawa. HRK: Hadley R. Kutcher. VK: Valentyna Klymiuk. GB: Guus Bakkeren. CP: Curtis Pozniak. GSB: Gurcharn S. Brar

**Project conception & hypotheses**: **GSB**

**Project administration and supervision:** GSB

**Grant writing and supervision**: GSB, GB, HRK, CP

**Experimental design**: SH, ML, CP, GSB, SF

**Manuscript writing**: SH, ML, GSB

**Manuscript revisions**: All Authors

**Isolate handling and phenotyping**: MA, RB, UG, AF, GSB

**Isolate collection**: MA, RB, AF, USG, BM, HSR, HRK, VK, GSB

**DNA extraction, purification, and library preparation**: ML, SHK, JB

**Genome sequencing and assembly**: JB, SW, SH

**Genome annotation:** SH

**Genome analysis**: SH, ML

## Supplementary files

**Supplementary file S1**. Distance matrix between haplotype genomes. Each phased haploid genome was aligned to each other, and the percent coverage (lower half) and SNPs/kbp (upper half) were assessed. Notable SNPs/kbp divide well into two categories: low SNPs/kbp (green) and high (yellow), which are indicative of whether the haploid genomes are within the same overall Group. Note the extremely close alignment between Yr2016-44 A and W047-1 A.

**Supplementary file S2.** Singleton gene clusters used in Anvi’o analysis and their functional annotations, and distributions across each haplotype genome.

**Supplementary file S3**. DNA integrity test of HMW Pst and Psh DNA samples on TapeStation.

**Supplementary file S4**. 16,145 pan-genomic clusters of 54 *Pst* and *Psh* haplotypic, scaffold-phased and annotated genomes are aligned on the chromosome location. The genomes are organized in radial bins of the outside layer as 18 chromosomes (Chr_1 to Chr_18) and unassigned locations (ChrUn). Genomes of haplotype A, B, C, D, E, F, G, H, I, J and K are arranged in the order generated by phylogenetic tree of average nucleotide identity (ANI) and colored by red, blue, yellow, dark gray, light gray, purple, light green, yellow and cyan, respectively. The detailed location, sequences and functions of each gene clusters per haploid genome is described in Supplementary file 9, 10 and 11.

**Supplementary file S5.** Core and single-copy coregene clusters used in Anvi’o analysis and their functional annotations, and distributions across each haplotype genome.

**Supplementary file S6.** Accessory gene clusters used in Anvi’o analysis and their functional annotations, and distributions across each haplotype genome.

**Supplementary table 7.** Functional roles of haplotype-associated Pfam domains and COG20-group proteins in fungal virulence and host adaption. The gene clusters enriched in more than three haplotypes were selected from Accessory gene clusters used in Anvi’o pan-genome and generated by frequency table of functional enrichment test implemented by Amy Willis in R (https://github.com/adw96), and filtered by using a Generalized Linear Model with the logit linkage to set a threshold of enrichment score > 20 and p/q-value < 0.05 for each function truly enriched in certain groups.

## References

1. Chen, X. M. Epidemiology and control of stripe rust *Puccinia striiformis* f. sp. *tritici* on wheat. Canadian Journal of Plant Pathology. 27(3):314–37. (2005)

2. Watson, I. A. Wheat and its rust parasites in Australia. Wheat Science-Today and Tomorrow, 129–147 (1981)

3. McIntosh, R. A. Wellings, C. R., & Park, R. F. Wheat rusts: an atlas of resistance genes. CSIRO publishing (1995).

4. Li, F., Upadhyaya, N.M., Sperschneider, J., Matny, O., Nguyen-Phuc, H., Mago, R., Raley, C., Miller, M.E., Silverstein, K.A., Henningsen, E. and Hirsch, C.D. Emergence of the Ug99 lineage of the wheat stem rust pathogen through somatic hybridisation. Nature Communications, 10(1) 5068 (2019)

5. Schwessinger, B., Chen, Y.J., Tien, R., Vogt, J.K., Sperschneider, J., Nagar, R., McMullan, M., Sicheritz-Ponten, T., Sørensen, C.K., Hovmøller, M.S. and Rathjen, J.P. Distinct life histories impact dikaryotic genome evolution in the rust fungus *Puccinia striiformis* causing stripe rust in wheat. Genome biology and evolution, 12(5) 597–617 (2020)

6. Goddard, M.V. The production of a new race, 105 E 137 of Puccinia striiformis in glasshouse experiments. Transactions of the British Mycological Society, 67(3), pp.395–398. (1976)

7. Park, R. F., & Wellings, C. R. Somatic hybridization in the Uredinales. Annual review of Phytopathology, 50(1), 219–239 (2012)

8. Watson, I. A. Wheat and its rust parasites in Australia. Wheat science-today and tomorrow, 129–147. (1981)

9. Brar, G.S., Fetch, T., McCallum, B.D., Hucl, P.J. and Kutcher, H.R. Virulence dynamics and breeding for resistance to stripe, stem, and leaf rust in Canada since 2000. Plant disease, 103(12), pp.2981–2995 (2019)

10. Wang, M. N., & Chen, X. M. Barberry does not function as an alternate host for *Puccinia striiformis* f. sp. *tritici* in the US Pacific Northwest due to teliospore degradation and barberry phenology. Plant Disease, 99(11), 1500–1506 (2015)

11. Chen, X., Penman, L., Wan, A., & Cheng, P. Virulence races of *Puccinia striiformis* f. sp. *tritici* in 2006 and 2007 and development of wheat stripe rust and distributions, dynamics, and evolutionary relationships of races from 2000 to 2007 in the United States. Canadian Journal of Plant Pathology, 32(3), 315–333. (2010)

12. Brar, G.S., Ali, S., Qutob, D., Ambrose, S., Lou, K., Maclachlan, R., Pozniak, C.J., Fu, Y.-B., Sharpe, A.G. and Kutcher, H.R. (2018), Genome re-sequencing and simple sequence repeat markers reveal the existence of divergent lineages in the Canadian Puccinia striiformis f. sp. tritici population with extensive DNA methylation. Environ Microbiol, 20: 1498–1515.

13. Brar, G. S., & Kutcher, H. R. Race characterization of *Puccinia striiformis* f. sp. *tritici*, the cause of wheat stripe rust, in Saskatchewan and Southern Alberta, Canada and virulence comparison with races from the United States. Plant Disease, 100(8), 1744–1753. (2016)

14. Ghanbarnia, K., Gourlie, R., Amundsen, E., Aboukhaddour, R. The changing virulence of stripe rust in Canada from 1984 to 2017. Phytopathology, 111(10), 1840–1850. (2021)

15 Milus E. A., Kristensen K., Hovmøller M. S., Evidence for Increased Aggressiveness in a Recent Widespread Strain of *Puccinia striiformis* f. sp. *tritici* Causing Stripe Rust of Wheat. Phytopathology, 99(1) (2009)

16. Holden S., Bakkeren G., Hubensky J., Bamrah R., Abbasi M., Qutob D., de Graaf M. L., Kim S. H., Kutcher H. R., McCallum B. D., Randhawa H. S. Uncovering the history of recombination and population structure in western Canadian stripe rust populations through mating type alleles. BMC biology. 21(1), 233 (2023)

17. Schwessinger, B., Sperschneider, J., Cuddy, W.S., Garnica, D.P., Miller, M.E., Taylor, J.M., Dodds, P.N., Figueroa, M., Park, R.F. and Rathjen, J.P. A near-complete haplotype-phased genome of the dikaryotic wheat stripe rust fungus *Puccinia striiformis* f. sp. *tritici* reveals high inter-haplotype diversity. MBio, 9(1) 10–1128 (2018)

18. Schwessinger, B., Jones, A., Albekaa, M., Hu, Y., Mackenzie, A., Tam, R., Nagar, R., Milgate, A., Rathjen, J.P. and Periyannan, S. A Chromosome Scale Assembly of an Australian *Puccinia striiformis* f. sp. *tritici* Isolate of the PstS1 Lineage. Molecular Plant-Microbe Interactions, 35(3), pp.293–296. (2022)

19. Wang, J., Xu, Y., Peng, Y., Wang, Y., Kang, Z. and Zhao, J. A fully haplotype-resolved and nearly gap-free genome assembly of wheat stripe rust fungus. Scientific Data, 11(1), 508 (2024)

20. Miller, M.E., Zhang, Y., Omidvar, V., Sperschneider, J., Schwessinger, B., Raley, C., Palmer, J.M., Garnica, D., Upadhyaya, N., Rathjen, J. and Taylor, J.M. De novo assembly and phasing of dikaryotic genomes from two isolates of *Puccinia coronata* f. sp. *avenae*, the causal agent of oat crown rust. MBio, 9(1), 10–1128 2018

21. Hewitt, T.C., Henningsen, E.C., Pereira, D., McElroy, K., Nazareno, E.S., Dugyala, S., Nguyen-Phuc, H., Li, F., Miller, M.E., Visser, B. and Pretorius, Z.A. Genome-enabled analysis of population dynamics and virulence-associated loci in the oat crown rust fungus *Puccinia coronata* f. sp. *avenae*. Molecular Plant-Microbe Interactions, 37(3), 290–303 (2024)

22. Sperschneider, J., Hewitt, T., Lewis, D. C., Periyannan, S., Milgate, A. W., Hickey, L. T., Mago R., Dodds P. N., Figueroa, M. Nuclear exchange generates population diversity in the wheat leaf rust pathogen *Puccinia triticina*. Nature Microbiology, 8(11), 2130–2141 (2023)

23. Vasquez-Gross, H., Kaur, S., Epstein, L. and Dubcovsky, J. A haplotype-phased genome of wheat stripe rust pathogen *Puccinia striiformis* f. sp. *tritici*, race PST-130 from the Western USA. PLoS One, 15(11), 0238611 (2020)

24. Wang, Y., Yin, M., & He, F. Two divergent haploid nuclei shaped the landscape of population diversity in wheat stripe rust. bioRxiv, 2024-12. (2024)

25. Duplessis, S., Cuomo, C.A., Lin, Y.C., Aerts, A., Tisserant, E., Veneault-Fourrey, C., Joly, D.L., Hacquard, S., Amselem, J., Cantarel, B.L. and Chiu, R., Obligate biotrophy features unraveled by the genomic analysis of rust fungi. Proceedings of the National Academy of Sciences, 108(22), 9166– 9171. (2011)

26. Soyer, J.L., El Ghalid, M., Glaser, N., Ollivier, B., Linglin, J., Grandaubert, J., Balesdent, M.H., Connolly, L.R., Freitag, M., Rouxel, T. and Fudal, I. Epigenetic control of effector gene expression in the plant pathogenic fungus Leptosphaeria maculans. PLoS Genetics, 10(3), e1004227. (2014)

27. Muszewska, A., Hoffman-Sommer, M., Grynberg, M. Retrotransposons in fungi: drivers of genome evolution. PLoS ONE, 6(6), e19387 (2011).

28. Spanu, P.D., Abbott, J.C., Amselem, J., Burgis, T.A., Soanes, D.M., Stüber, K., Loren van Themaat, E.V., Brown, J.K., Butcher, S.A., Gurr, S.J. and Lebrun, M.H. Genome expansion and gene loss in powdery mildew fungi reveal trade offs in extreme parasitism. Science, 330(6010), 1543–1546. (2010)

29. Raffaele, S., Kamoun, S. Genome evolution in filamentous plant pathogens: why bigger can be better. Nature Reviews Microbiology, 10, 417–430. (2012)

30. Makarova, K.S., Aravind, L. and Koonin, E.V., 2002. SWIM, a novel Zn-chelating domain present in bacteria, archaea and eukaryotes. Parks, L. W., et al. Regulation of sterol biosynthesis in yeast. Lipids, 34(3), 299–309. (1999)

31. Hirmondo, R., Lopata, A., Suranyi, E.V., Vertessy, B.G., Toth, J. Differential control of dNTP biosynthesis and genome integrity maintenance by the dUTPase superfamily enzymes. Scientific reports, 7(1), p.6043. (2017)

32. Mora-Montes, H.M., Ponce-Noyola, P., Villagómez-Castro, J.C., Gow, N.A., Flores-Carreón, A., López-Romero, E. Protein glycosylation in Candida. Future microbiology, 4(9), pp.1167–1183. (2009)

33. Steenwyk, J.L. Evolutionary divergence in DNA damage responses among fungi. MBio, 12(2), pp.10–1128. (2021)

34. Helmlinger, D. and Tora, L. Sharing the SAGA. Trends in biochemical sciences, 42(11), pp.850–861. (2017)

35. Crunden, J.L. Diezmann, S. Hsp90 interaction networks in fungi—tools and techniques. FEMS Yeast Research, 21(7), p.foab054. (2021)

36. Gerwien, F., Skrahina, V., Kasper, L., Hube, B., Brunke, S. Metals in fungal virulence. FEMS microbiology reviews, 42(1), p.fux050 (2018)

37. Pritchard, L., Glover, R. H., Humphris, S., Elphinstone, J. G., & Toth, I. K. Genomics and taxonomy in diagnostics for food security: soft-rotting enterobacterial plant pathogens. Analytical methods, 8(1), 12–24. (2016)

38. Kulshrestha, A., & Gupta, P. Secreted Aspartyl Proteases Family: A Perspective Review on the Regulation of Fungal Pathogenesis. Future Microbiology, 18(5), 295–309. (2023) 10.2217/fmb-2022-0143

39. Li, Y., Kabbage, M., Liu, W., & Dickman, M. B. Aspartyl Protease-Mediated Cleavage of BAG6 Is Necessary for Autophagy and Fungal Resistance in Plants. The Plant cell, 28(1), 233–247. (2016) 10.1105/tpc.15.00626

40. Zawrotniak, M., Satala, D., Juszczak, M. Bras, G., Rapala-Kozik, M. Candida albicans aspartyl protease (Sap6) inhibits neutrophil function via a “Trojan horse” mechanism. Sci Rep 15, 6946 (2025). 10.1038/s41598-025-91425-x

41. Haigh, J. The accumulation of deleterious genes in a population—Muller’s ratchet. Theoretical population biology, 14(2), pp.251–267 (1978)

42. Wan, A., Chen, X., & Yuen, J. Races of Puccinia striiformis f. sp. tritici in the United States in 2011 and 2012 and comparison with races in 2010. Plant disease, 100(5), 966–975 (2016)

43. Li, M., Brar, G. S., Schwessinger, B., & Jones, A. High Molecular Weight DNA Extraction from Wheat Rust Spores. In Wheat Rusts and Resistance Breeding: Methods and Protocols 139–149. Springer US. (2025)

44. Andrews, S. FastQC: A Quality Control Tool for High Throughput Sequence Data [Online Resource]. (2010) http://www.bioinformatics.babraham.ac.uk/projects/fastqc/

45. Mapleson, D., Garcia Accinelli, G., Kettleborough, G., Wright, J., & Clavijo, B. J. KAT: a K-mer analysis toolkit to quality control NGS datasets and genome assemblies. Bioinformatics, 33(4), 574–576. (2017)

46. Cheng, H., Concepcion, G. T., Feng, X., Zhang, H., & Li, H. Haplotype-resolved de novo assembly using phased assembly graphs with hifiasm. Nature methods, 18(2), 170–175 (2021)

47. Li, H. Minimap2: pairwise alignment for nucleotide sequences. Bioinformatics, 34(18), 3094–3100. (2018)

48. Delcher, A. L., Salzberg, S. L., & Phillippy, A. M. Using MUMmer to identify similar regions in large sequence sets. Current protocols in bioinformatics, (1), 10–3. (2003)

49. Alonge, M., Lebeigle, L., Kirsche, M., Jenike, K., Ou, S., Aganezov, S., Wang, X., Lippman, Z.B., Schatz, M.C. and Soyk, S. Automated assembly scaffolding using RagTag elevates a new tomato system for high-throughput genome editing. Genome biology, 23(1) 258 (2022)

50. Jonathan M. Palmer, & Jason Stajich. Funannotate v1.8.1: Eukaryotic genome annotation (v1.8). (2020) Zenodo. 10.5281/zenodo.1134477

51. Ou S., Su W., Liao Y., Chougule K., Agda J. R. A., Hellinga A. J., Lugo C. S. B., Elliott T. A., Ware D., Peterson T., Jiaradhng N., Hirsch C. N., Hufford M. B. Benchmarking Transposable Element Annotation Methods for Creation of a Streamlined, Comprehensive Pipeline. Genome Biol. 20(1): 275. (2019)

52. Radhakrishnan, G.V., Cook, N.M., Bueno-Sancho, V., Lewis, C.M., Persoons, A., Mitiku, A.D., Heaton, M., Davey, P.E., Abeyo, B., Alemayehu, Y., Badebo, A., MARPLE, a point-of-care, strain-level disease diagnostics and surveillance tool for complex fungal pathogens. BMC biology, 17(1), 65. (2019)

53. Eren, A. M., Esen, Ö. C., Quince, C., Vineis, J. H., Morrison, H. G., Sogin, M. L., Delmont, T. O. Anvi’o: an advanced analysis and visualization platform for ‘omics data. PeerJ, 3, e1319. (2015)

