## Supplementary File S2 for "Wheat and barley stripe rust pan-genome facilitates discovery of the predominant North American lineage’s origin in somatic hybridization"

### Slide 1
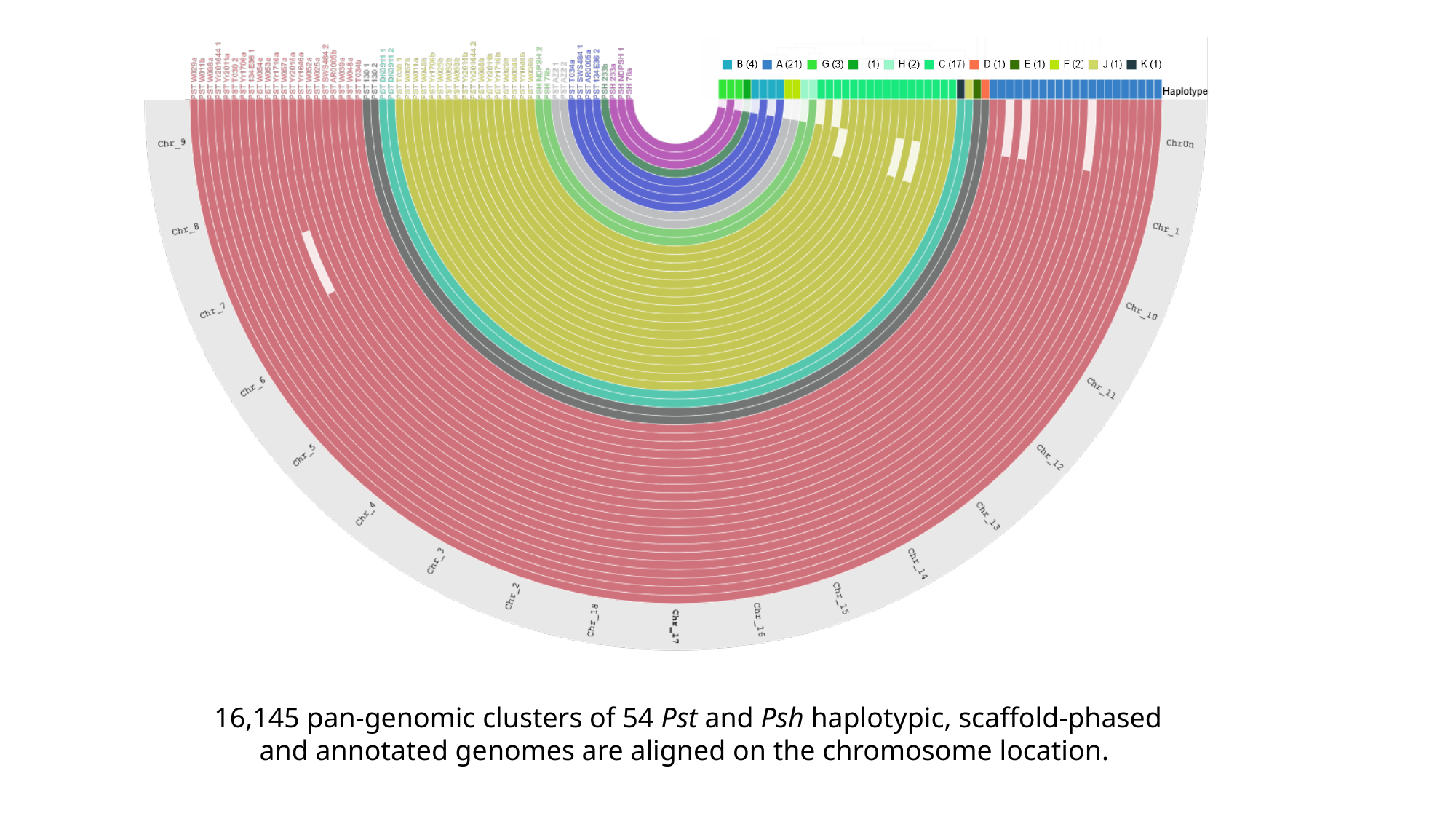

16,145 pan-genomic clusters of 54 Pst and Psh haplotypic, scaffold-phased and annotated genomes are aligned on the chromosome location.
