## Supplementary File S3 for "Wheat and barley stripe rust pan-genome facilitates discovery of the predominant North American lineage’s origin in somatic hybridization"

### Slide 1
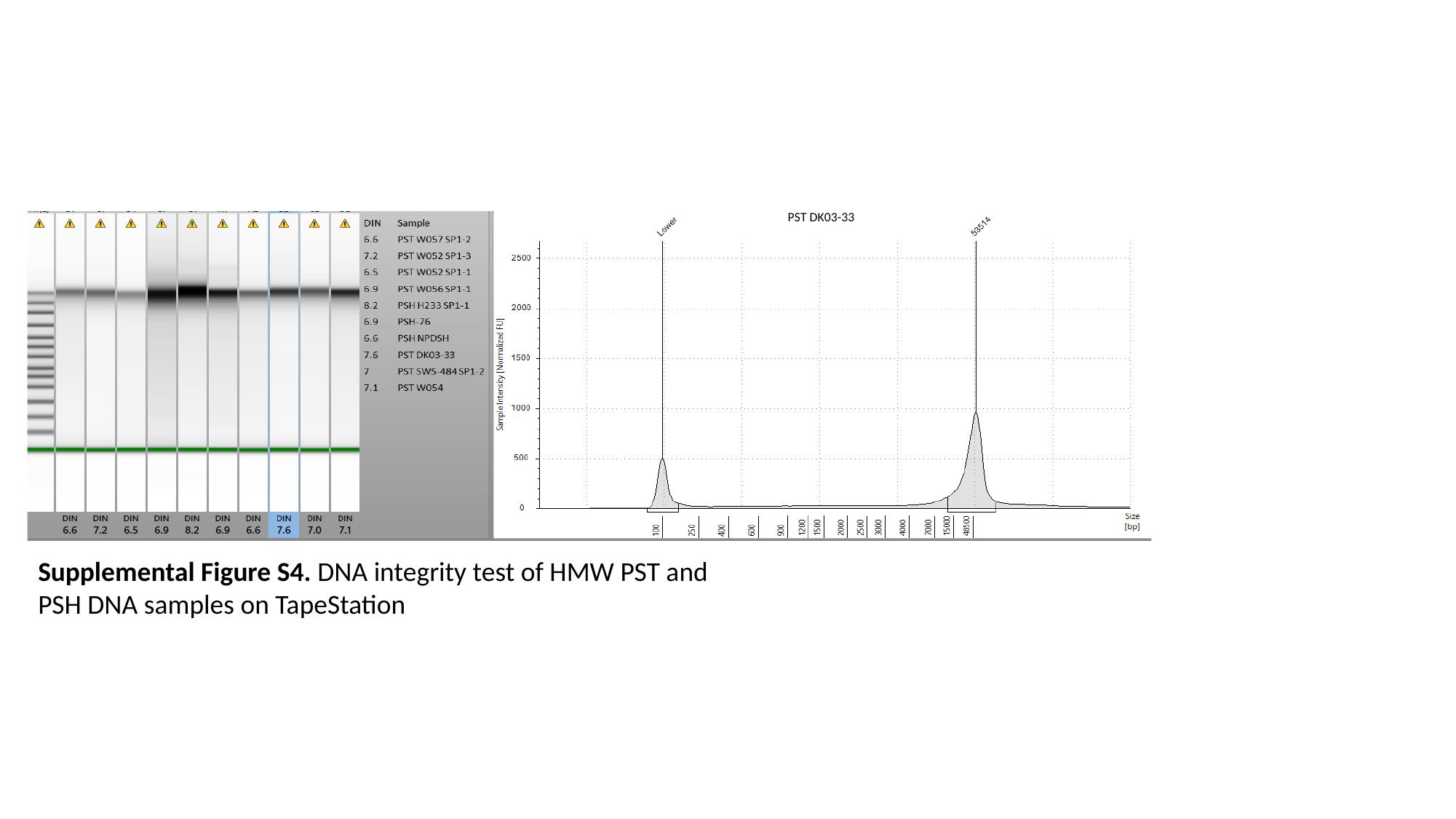

PST DK03-33
Supplemental Figure S4. DNA integrity test of HMW PST and PSH DNA samples on TapeStation
