## Supplementary File S1 for "Wheat and barley stripe rust pan-genome facilitates discovery of the predominant North American lineage’s origin in somatic hybridization"

### Slide 1
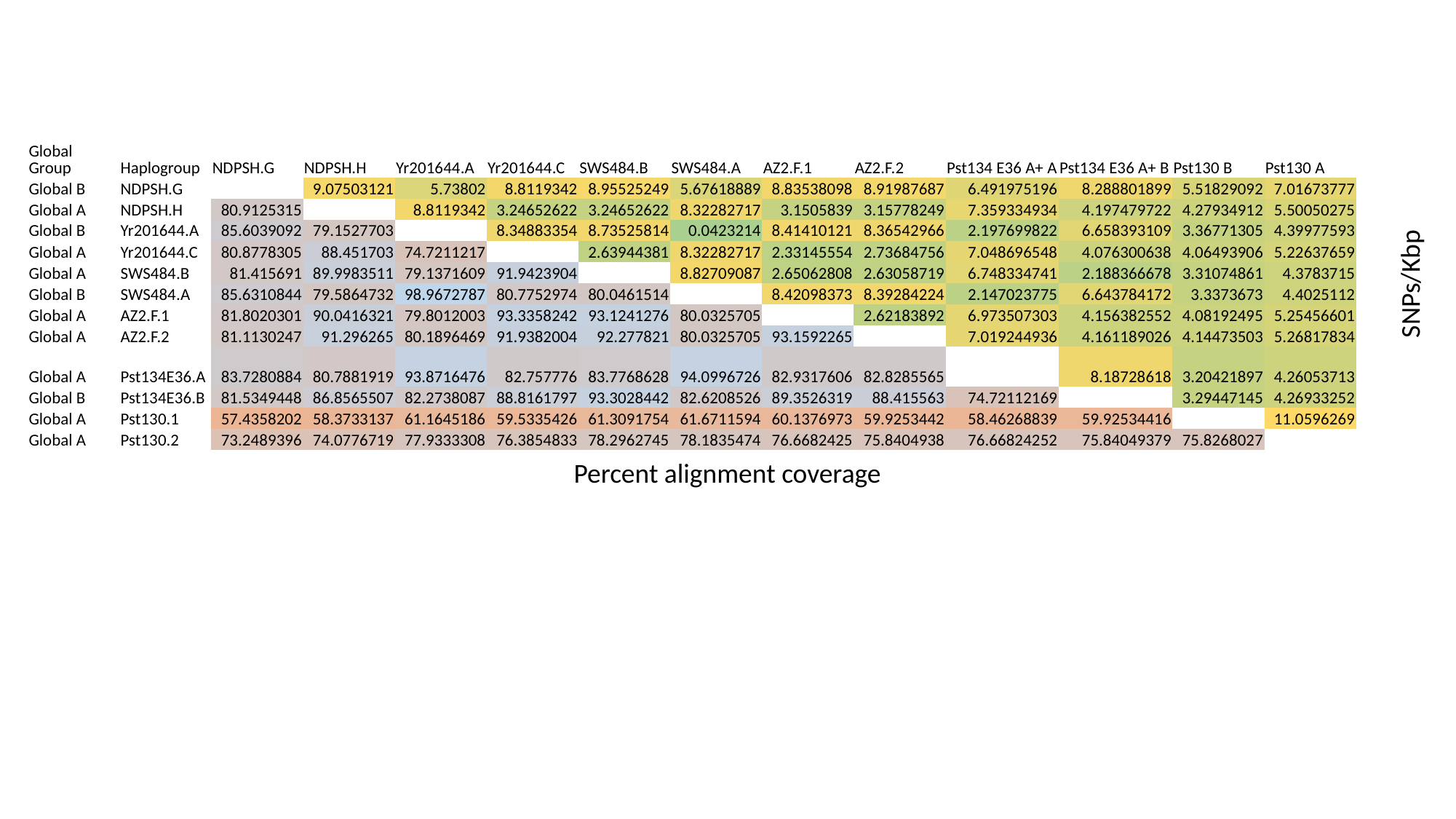

| Global Group | Haplogroup | NDPSH.G | NDPSH.H | Yr201644.A | Yr201644.C | SWS484.B | SWS484.A | AZ2.F.1 | AZ2.F.2 | Pst134 E36 A+ A | Pst134 E36 A+ B | Pst130 B | Pst130 A |
| --- | --- | --- | --- | --- | --- | --- | --- | --- | --- | --- | --- | --- | --- |
| Global B | NDPSH.G | | 9.07503121 | 5.73802 | 8.8119342 | 8.95525249 | 5.67618889 | 8.83538098 | 8.91987687 | 6.491975196 | 8.288801899 | 5.51829092 | 7.01673777 |
| Global A | NDPSH.H | 80.9125315 | | 8.8119342 | 3.24652622 | 3.24652622 | 8.32282717 | 3.1505839 | 3.15778249 | 7.359334934 | 4.197479722 | 4.27934912 | 5.50050275 |
| Global B | Yr201644.A | 85.6039092 | 79.1527703 | | 8.34883354 | 8.73525814 | 0.0423214 | 8.41410121 | 8.36542966 | 2.197699822 | 6.658393109 | 3.36771305 | 4.39977593 |
| Global A | Yr201644.C | 80.8778305 | 88.451703 | 74.7211217 | | 2.63944381 | 8.32282717 | 2.33145554 | 2.73684756 | 7.048696548 | 4.076300638 | 4.06493906 | 5.22637659 |
| Global A | SWS484.B | 81.415691 | 89.9983511 | 79.1371609 | 91.9423904 | | 8.82709087 | 2.65062808 | 2.63058719 | 6.748334741 | 2.188366678 | 3.31074861 | 4.3783715 |
| Global B | SWS484.A | 85.6310844 | 79.5864732 | 98.9672787 | 80.7752974 | 80.0461514 | | 8.42098373 | 8.39284224 | 2.147023775 | 6.643784172 | 3.3373673 | 4.4025112 |
| Global A | AZ2.F.1 | 81.8020301 | 90.0416321 | 79.8012003 | 93.3358242 | 93.1241276 | 80.0325705 | | 2.62183892 | 6.973507303 | 4.156382552 | 4.08192495 | 5.25456601 |
| Global A | AZ2.F.2 | 81.1130247 | 91.296265 | 80.1896469 | 91.9382004 | 92.277821 | 80.0325705 | 93.1592265 | | 7.019244936 | 4.161189026 | 4.14473503 | 5.26817834 |
| Global A | Pst134E36.A | 83.7280884 | 80.7881919 | 93.8716476 | 82.757776 | 83.7768628 | 94.0996726 | 82.9317606 | 82.8285565 | | 8.18728618 | 3.20421897 | 4.26053713 |
| Global B | Pst134E36.B | 81.5349448 | 86.8565507 | 82.2738087 | 88.8161797 | 93.3028442 | 82.6208526 | 89.3526319 | 88.415563 | 74.72112169 | | 3.29447145 | 4.26933252 |
| Global A | Pst130.1 | 57.4358202 | 58.3733137 | 61.1645186 | 59.5335426 | 61.3091754 | 61.6711594 | 60.1376973 | 59.9253442 | 58.46268839 | 59.92534416 | | 11.0596269 |
| Global A | Pst130.2 | 73.2489396 | 74.0776719 | 77.9333308 | 76.3854833 | 78.2962745 | 78.1835474 | 76.6682425 | 75.8404938 | 76.66824252 | 75.84049379 | 75.8268027 | |
SNPs/Kbp
Percent alignment coverage
